# Predicting Potential Spawning Areas: a novel framework for elasmobranch conservation and spatial management

**DOI:** 10.1101/2025.02.11.637670

**Authors:** David Ruiz-García, Claudio Barría, Ana I. Colmenero, Juan A. Raga, David March

## Abstract

Defining and protecting critical habitats for elasmobranchs (sharks and rays), such as spawning areas, is essential for mitigating anthropogenic pressures that threaten their populations, primarily driven by fisheries and habitat degradation. This study presents a novel modelling-based framework to identify Potential Spawning Areas (PSAs) - habitats offering optimal conditions for oviposition. Using fisheries-dependent trawl bycatch data combined with environmental and anthropogenic predictors, we applied machine-learning models to delineate PSAs for the smallspotted catshark (*Scyliorhinus canicula*) and skates (*Raja spp.*) in the western Mediterranean. Static environmental predictors, including depth and slope, were primary drivers, while dynamic predictors, including sea bottom temperature and salinity played seasonally relevant roles. Trawl fishing pressure also influenced importantly the distribution of PSAs, revealing a concerning positive feedback loop between exploitation and habitat degradation. While PSAs experienced lower fishing effort than the rest of the study area, a substantial proportion of egg case bycatch still occurs within them. The current network of Marine Protected Areas in the region fails to adequately safeguard these habitats due to limited coverage and enforcement. Our findings underscore significant gaps in spatial management and the urgent need for targeted conservation measures. The PSA framework provides a robust, scalable tool for identifying critical habitats across regions and species, offering actionable insights for marine spatial planning and ecosystem-based management. This adaptable approach can support global conservation efforts for elasmobranchs and their ecosystems.

**Significance:** Sharks and rays are vital to marine ecosystems but face alarming extinction risks due to overfishing and habitat loss. Identifying critical reproductive habitats, such as spawning areas, is essential for protection but remains a significant challenge. This study introduces a novel framework for identifying Potential Spawning Areas (PSAs) - habitats offering optimal conditions for oviposition - using machine-learning and fisheries bycatch data. Applied in the western Mediterranean for the smallspotted catshark and skates, it demonstrates how environmental factors and fishing pressures shape these habitats, highlighting significant management gaps. The PSAs framework offers a scalable approach to guide the establishment of marine protected areas and fisheries management plans, providing a novel approach for conserving vulnerable marine species and their habitats.

## INTRODUCTION

Identifying and delineating critical habitats is fundamental for designing effective area-based conservation strategies that mitigate anthropogenic pressures (1). This is particularly important for elasmobranchs (sharks and rays), which face increasing extinction risks globally (2). Elasmobranchs play a key role in the regulation of marine ecosystems by controlling the dynamics of their prey populations and connecting trophic webs across habitats (3). Their life history traits (slow growth, late maturity, low fecundity, and longevity) make them particularly vulnerable to anthropogenic pressures, primarily driven by fisheries and habitat degradation (2). Among elasmobranchs, oviparous species tend to have higher fecundity and maximum population growth rates than viviparous species, resulting in a lower vulnerability (4). However, oviparous species may be exposed to an overlooked interplay between fisheries exploitation and habitat degradation, which can directly and indirectly disrupt critical spawning areas, ultimately increasing their vulnerability (5, 6).

Nursery areas are critical habitats that support stock recruitment while generating cascading benefits that strengthen broader ecological functions and services (7). Specific criteria have been set to recognise nursery areas for elasmobranchs (8). Recently, the concept of nursery areas has been expanded by some authors to include areas where elasmobranch spawning preferentially occurs, defining them as “egg nurseries” (9–12). These areas are characterized by 1) a high density of egg cases, 2) egg cases in contact with fixed substrates (biogenic or geological), and 3) repeated use for oviposition over multiple years, although neonates and juveniles may migrate to different areas. A minimum threshold of 1,000 egg cases per km² has been established to identify areas of high-density from trawl surveys, with such accumulations typically confined to small areas (10, 12, 13).

Elasmobranch egg nurseries have been documented across different habitats, ranging from structured bottoms such as high-energy rocky areas, coral reefs, and algal beds, to low-energy soft bottoms composed by sandy and muddy substrates, frequently colonised by filter feeders such as sponges or cnidarians (10–16). These nurseries hold significant ecological importance and, when geographically restricted, they can be highly vulnerable to human disturbances (9). Both structured and soft sediment habitats hosting egg nurseries are particularly susceptible to anthropogenic disturbances. However, soft sediments are more frequently impacted by demersal trawling activities, constituting a primary driver of habitat degradation (17). Additionally, climate change and pollution have also emerged as important contributors for habitat degradation, potentially altering elasmobranch spawning patterns (6, 18, 19).

Knowledge of elasmobranch habitat use remains limited, with critical areas like nurseries remaining unidentified for most species (1, 20). The philopatric behaviour reported in nurseries underscores their importance in sustaining local populations across generations (8, 12, 21). Understanding the spatial distribution of these critical habitats is crucial for guiding fisheries management and marine spatial planning (9). Stakeholders, including the International Council for the Exploration of the Sea (ICES) and the General Fisheries Commission for the Mediterranean (GFCM), emphasize protecting elasmobranch spawning and nursery grounds, classifying them as Essential Fish Habitats (22–24). To further this effort, the International Union for the Conservation of Nature (IUCN) recently launched the project Important Shark and Ray Areas (ISRA). This initiative aims to identify critical areas for elasmobranchs and facilitate decision-making, with spawning and nursery areas composing one of the main criteria for their designation (1).

Defining elasmobranch egg nursery areas is challenging due to limited data, with only a few studies worldwide reporting egg case occurrence and abundance, mainly using trawl data and opportunistic observations from imaging systems (10, 15). Certainly, egg cases are infrequently recorded in fisheries-independent trawl surveys, and no reports exist for bycatch in commercial fisheries, underscoring a key knowledge gap regarding fisheries-related threats (9, 12). Additionally, imaging systems, such as ROVs, are rarely used beyond specific locations (13–16). Despite these challenges, oviposition patterns have been linked to environmental factors such as temperature, sediment type and currents (12, 20), alongside human impacts like bottom trawling effort and pollution (6, 10). Consequently, the spatial distribution of spawning areas is likely shaped by species-specific physiological tolerances to environmental gradients and the ecological status of the habitat, which can be influenced by anthropogenic pressures (17).

Given the challenges in identifying and mapping elasmobranch spawning areas, innovative approaches are needed to guide conservation efforts. We propose adopting the concept of *Potential Spawning Areas* (PSAs) - habitats shaped by the interplay of dynamic oceanographic and ecological factors alongside stable environmental features, converging to provide optimal conditions for spawning. Although this concept has not yet been applied to elasmobranchs, similar approaches have been successfully utilized for pelagic fish species whose eggs are part of the ichthyoplankton (25, 26). This approach acknowledges that PSAs may include regions where populations have declined, collapsed, or where access is restricted, meaning that spawning may not currently occur (26). Nonetheless, the identification of PSAs provides a valuable tool to prioritize research, inform conservation efforts or even guide reintroduction programs in such areas. Furthermore, we propose integrating PSAs extents across multiple species to define Aggregated Potential Spawning Areas (APSAs), representing regions of shared ecological importance for oviparous elasmobranchs.

In the western Mediterranean, where trawl fisheries have heavily impacted benthic habitats, elasmobranch spawning grounds and egg case bycatch rates remain poorly documented despite international efforts to define and protect them (27–29). To address this gap, we analyse fisheries-dependent trawl data, quantifying the commercial bycatch rates of elasmobranch egg cases and examining their spatial distribution patterns. We focus on the most frequently captured taxa, the smallspotted catshark (*Scyliorhinus canicula*), classified as *Least Concern* by the IUCN Red List, and skates of the genus *Raja*, encompassing at least five documented species, only one of which is classified as threatened by the same criteria. While these species are not highly threatened overall, monitoring the risks to their populations remains essential given their commercial interest, the absence of catch limits, shifting seafood markets, and the depletion of other target stocks in the study area, underscoring the need for proactive management (30).

Understanding bycatch dynamics in these taxa we aim to provide insights that inform conservation strategies for more vulnerable elasmobranchs while addressing key knowledge gaps regarding fisheries-related impacts on spawning habitats. Using a machine-learning modelling approach, we delineate PSAs for these taxa, assessing the effect of environmental and human drivers on their distribution. To further quantify fisheries impacts, we also calculate annual total bycatch estimates and integrate PSA predictions across species to define APSAs, assessing their overlap with trawl fishing effort and existing management measures. Through this approach, we aim to develop a scalable framework for identifying critical habitats, inform targeted conservation strategies and address gaps in marine spatial planning.

## RESULTS

### Egg case bycatch

To assess the distribution of spawning habitats for oviparous elasmobranch species, we analysed the occurrence and abundance of egg cases captured as bycatch in a commercial bottom trawl fishery (SI Appendix, Fig. S1). A total of 3,125 elasmobranch egg cases were found in 117 tows (60.3% of 194 analysed). Egg cases from at least seven species were recorded, including one shark and six skates (Table 1 and SI Appendix, Table S1). The smallspotted catshark egg cases were the most frequent, accounting also with the highest bycatch per unit of effort (BPUE) at a specific location, surpassing 1,000 egg cases per km^2^ (Table 1; Fig. 1). The skates included at least five species of the genus *Raja* and the longnosed skate (*Dipturus oxyrinchus*), the latter captured at very low frequency, near a submarine canyon (Table 1, SI Appendix, Fig. S2). Only 25.9% of *Raja* spp. egg cases were identified to the species level; therefore, data were aggregated at the genus level (see species-specific data in SI Appendix, Table S1). Hereafter, we refer to *Raja* spp. as skates, excluding *Dipturus oxyrinchus*. Skate egg cases were widespread across geographic and bathymetric ranges, with notable BPUE peaks in certain locations, approaching 1,000 egg cases per km^2^ (Table 1; Fig. 1).

**Fig. 1.**
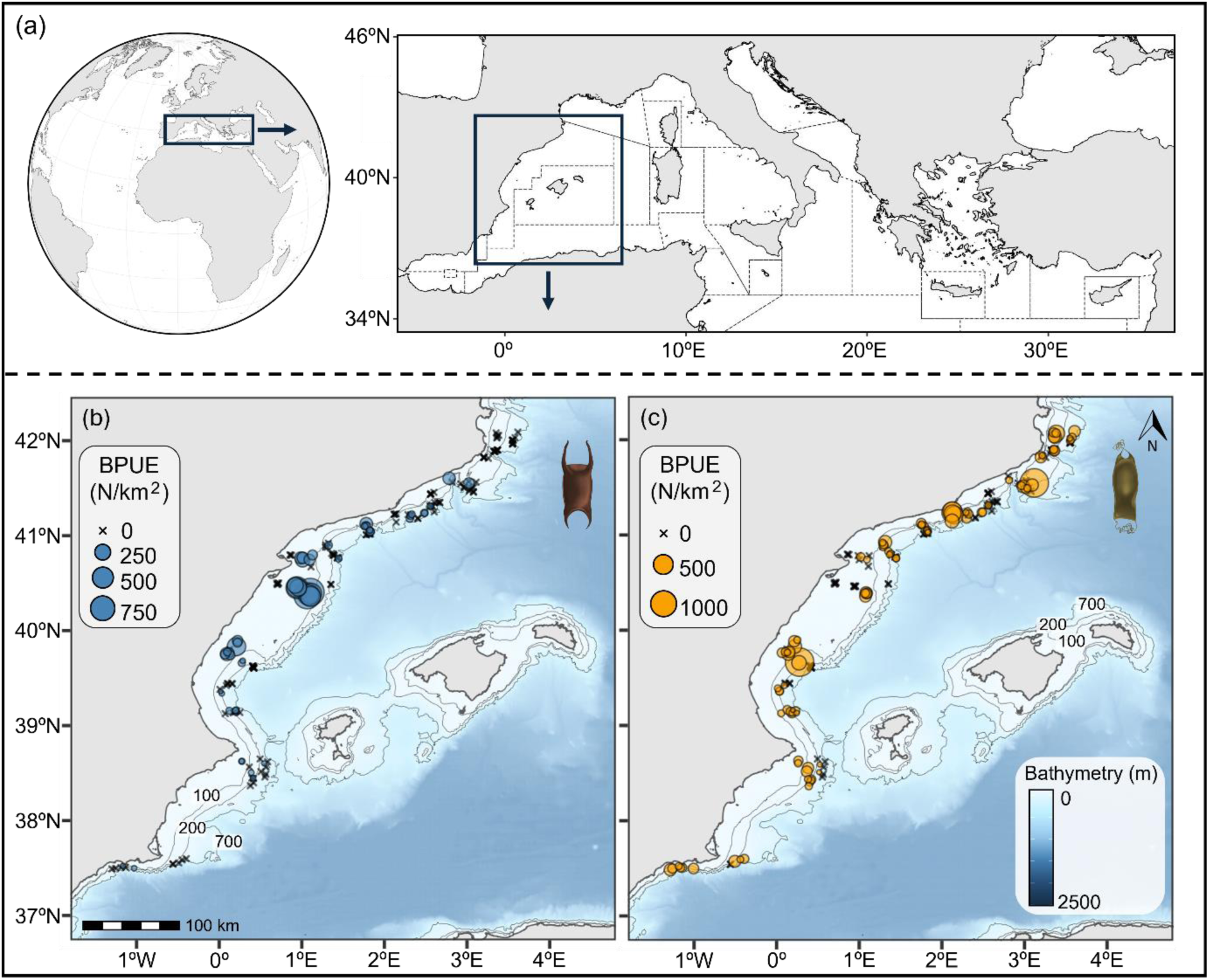
(a) Location of study area at a global scale and within the Mediterranean, accounting with GFCM geographical subareas; (b) Distribution of skate egg case BPUE, represented in blue; and (c) smallspotted catshark egg case BPUE, represented in orange. Circles represent egg case presence, with size proportional to BPUE, while crosses denote their absence at sampling location.

**Table 1.** List of elasmobranch species for which egg cases were captured, with their conservation status based on the regional assessment of the Mediterranean IUCN Red List. Reported values include the total number of egg cases captured (N_T_), the maximum BPUE (N/km^2^) with the corresponding number of egg cases (n) in a single tow, the mean BPUE (N/km^2^), the frequency of occurrence (Freq.) across all surveyed tows, and the depth range in which they occurred.

| Family | Species | IUCN status | $N_T$ | Maximum | | Mean BPUE | Freq. (%) | Depth (m) |
| --- | --- | --- | --- | --- | --- | --- | --- | --- |
|  |  |  |  | BPUE | n |  |  |  |
| Scyliorhinidae | *Scyliorhinus canicula* | LC | 2241 | 1469.3 | 600 | 53.4 ± 168.3 | 52.6 | 55-600 |
| Rajidae | *Raja* spp. | - | 881 | 933.5 | 110 | 33.5 ± 117.7 | 30.9 | 55-600 |
|  | *Dipturus oxyrinchus* | NT | 3 | 17.3 | 1 | 0.2 ± 1.4 | 1.5 | 565-668 |

#### Effect of environmental and human drivers

To investigate the factors influencing the distribution and abundance of elasmobranch egg case bycatch, we analysed the effects of environmental and human drivers using Boosted Regression Trees (BRT) within a delta model framework. The evaluation of the delta model components (occurrence and abundance) through cross-validation procedures indicated a good fit to the observed data. Both occurrence (presence/absence) models yielded AUC values of 0.8 and explained a 29.5% and 19.8% of the cross-validation deviance, while abundance (BPUE) models explained a 46.2% and 16.8% of the cross-validation deviance, for skates and the smallspotted catshark, respectively in both cases (SI Appendix, Tables S2-5).

The models revelated that the top three most influential drivers (i.e., fishing effort, depth and slope) were consistent across occurrence and abundance models, as well as between taxa, while some differences emerged in the shapes of their dependence curves (Fig. 2). Generally, bycatch occurrence and abundance decreased progressively with increasing fishing effort, except for the occurrence of smallspotted catshark egg cases, which remained consistent at moderate fishing effort levels before declining in highly trawled areas. In terms of depth, skate egg cases exhibited higher occurrence and abundance on the continental shelf, peaking between 50 and 100 m before progressively declining. In contrast, the smallspotted catshark egg case bycatch was greater along the shelf margin and upper slope, with a peak between 100 and 300 m, followed by a gradual decrease. A higher occurrence and density for skate egg cases was more likely in flat areas (near 0°), whereas for the smallspotted catshark, it was more likely in gently sloping areas (3°–8°; Fig. 2). Sea bottom salinity also played an influential role in predicting egg case occurrence for skates and abundance for the smallspotted catshark, both with higher rates in areas with slightly reduced sea salinity compared to the studied range (<38.25 psu). Additionally, sea bottom temperature also influenced egg case occurrence and abundance for the small spotted catshark, both with higher rates in temperatures ranging from 13°C to 14 °C (Fig. 2).

**Fig. 2.**
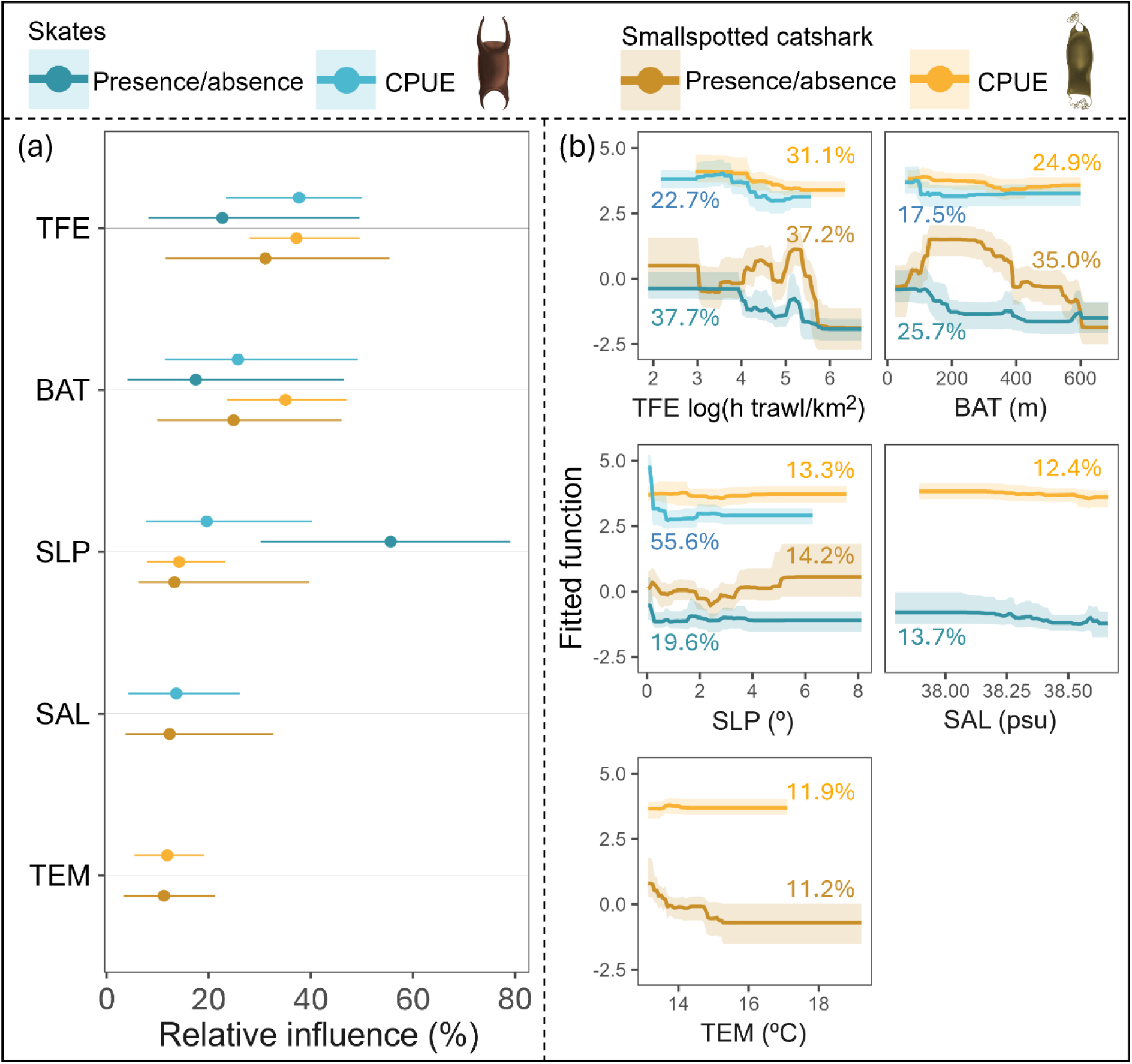
Effect of the environmental and human pressure drivers on egg case bycatch occurrence and log-transformed BPUE (log(x+1)) for skates and the smallspotted catshark, in blue and orange, respectively; (a) Relative influence of each predictor used to model bycatch occurrence and abundance. Dots represent medians and lines represent the 95% confidence interval range of the bootstrap predictions (n = 100); (b) Partial dependence plots of the predictors included in the BRT models. The mean relative influence in percentage is provided for each variable. The shading shows the 95% confidence interval estimated from 100 bootstrap samples of the data set. Predictor acronyms and definitions are described in Table 2.

**Table 2.**
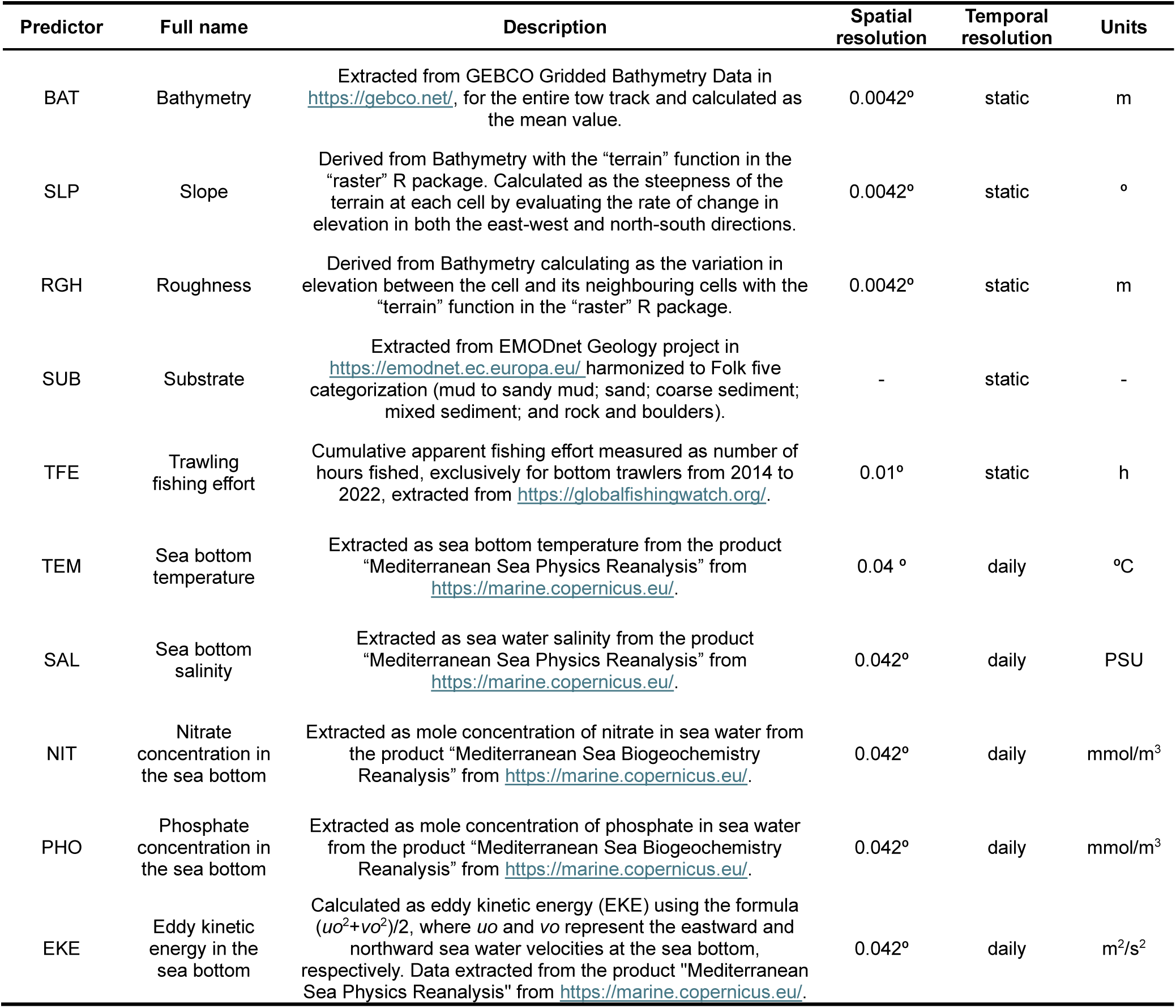
Predictor variables considered into the modelling approach.

#### Potential Spawning Areas

To identify Potential Spawning Areas (PSAs) for both species, we integrated daily occurrence and abundance predictions by computing their product. We then calculated seasonal and annual medians of the resulting egg case BPUE spatial predictions, along with the associated uncertainty, measured as the 95% confidence interval (CI). Areas within the upper decile of predicted median BPUE values were designated as PSAs, covering a total area of 3,478 km^2^. The primary PSAs for both species were identified along the extended continental shelf influenced by the Ebro River, spanning latitudes from 41.0° to 39.5° N (Fig. 3). Notably, the Intersection over Union (IoU) index revealed that annual PSAs of both species overlapped by 17.12%, with the entire overlap confined within these latitudinal bounds.

**Fig. 3.**
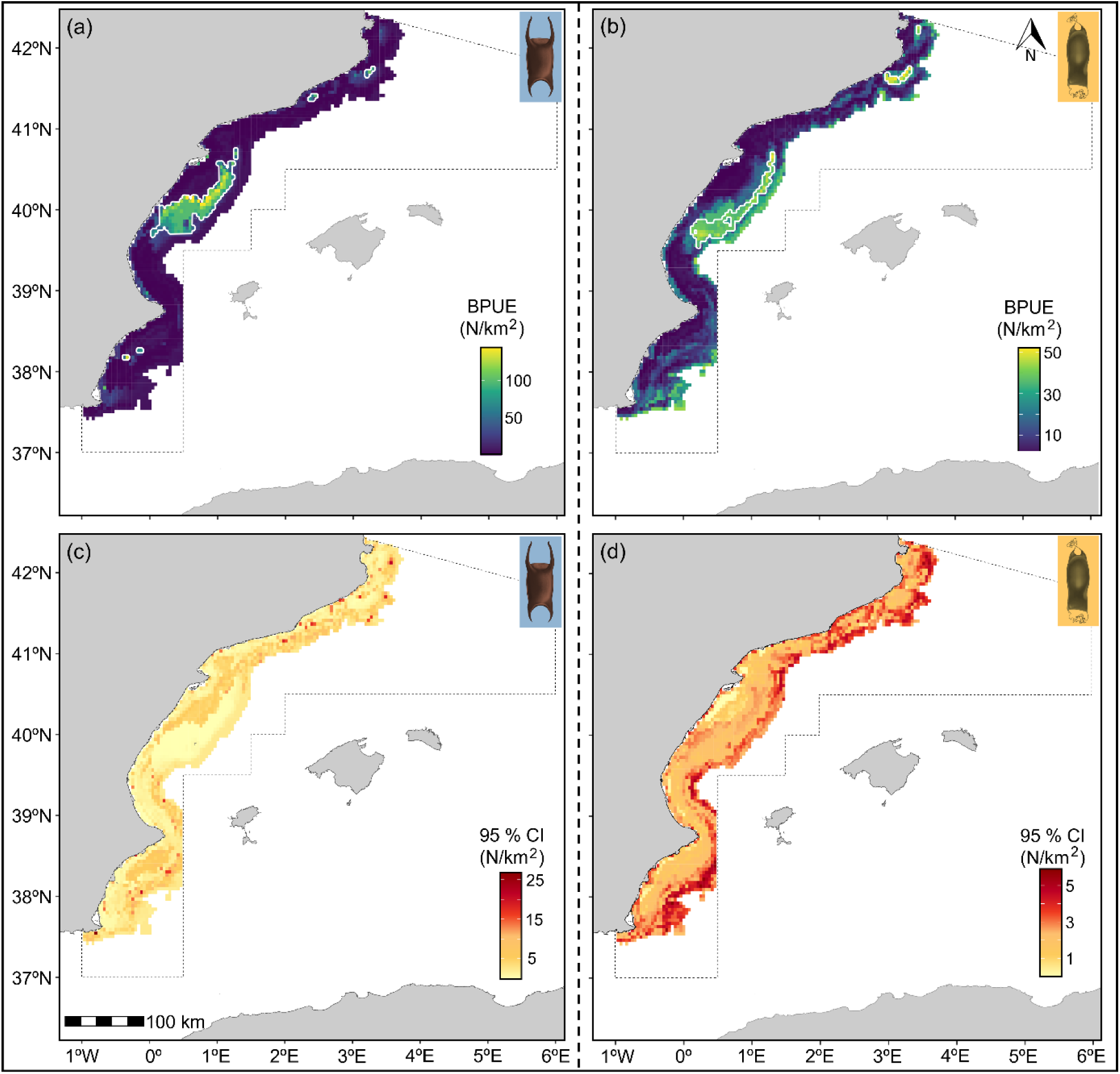
Spatial distribution of potential spawning areas and associated uncertainty. Annual median egg case BPUE predictions, based on daily medians of bootstrap models (n = 100), for skate species (a); and the smallspotted catshark (b). Uncertainty estimates, represented as the 95% confidence interval ranges, also derived from the daily medians of bootstrap models (n = 100), for skate species (c), and the smallspotted catshark (d). Potential spawning areas, determined as the upper decile of predicted BPUE, are highlighted within white contours.

#### Seasonal PSAs persistency

Seasonal persistence in the spatial distribution of PSAs (i.e., the proportion of seasons a cell was identified as part of a PSA) and variations in mean BPUE across the study area were analysed to assess the stability and consistency of these critical habitats. For skates, a high seasonal persistence was observed, along with non-significant variations in mean BPUE (Kruskal-Wallis test: χ^2^ = 0.67, df = 3, p = 0.88; SI Appendix, Supplementary Fig. S3,5). In contrast, the smallspotted catshark exhibited significant seasonal variation in mean BPUE across the study area (χ^2^ = 244.96, df = 3, p < 0.001). Pairwise Wilcoxon tests with Bonferroni correction identified two distinct seasonal groups: spring-winter and fall-summer. Seasonal persistence was high within the central areas of the seasonal PSAs, with some expansion into shallower regions during spring and winter (SI Appendix, Supplementary Fig. S4,5).

#### Estimation of total egg case bycatch

To further assess the impact of trawl bycatch on PSAs, we estimated the total number of egg cases accidentally captured in each bottom trawling metier operating in the study area during the predicted year (i.e., 2021). This estimation was based on the overlap between our daily median BPUE predictions (with associated 95% CI) and daily trawl fishing effort maps. Total effort amounted a total of 372,187 h, resulting in an estimated trawled area of 33,349 km^2^ across the study area in the year 2021. Total bycatch estimates indicate a capture of 233,239 ± 43,169 for skate egg cases and 228,935 ± 25,570 for smallspotted catshark egg cases, expressed as median and 95% CI, with the hake metier having the highest amount of egg case bycatch in both cases. A 42.0% and 8.3% of the total egg bycatch occurs within the PSAs for skates and the smallspotted catshark, respectively, while these areas cover 10% of the total study area (SI Appendix, Fig. S6).

#### Fishing pressure and spatial protection on APSAs

The annual PSAs for skates and the smallspotted catshark were integrated into Aggregated Potential Spawning Areas (APSAs), representing the combined spatial extent of critical oviposition habitats for the studied taxa. To assess anthropogenic pressure within these critical habitats, we compared trawl fishing effort intensity within APSAs to the rest of the study area. To evaluate protection levels, we quantified the overlap using the Intersection over Union index for existing management areas, including Marine Protected Areas (MPAs) and Trawling Exclusion Zones (TEZs). Additionally, we assessed overlap with Important Shark and Ray Areas (ISRAs), which, while designated for elasmobranch conservation, currently lack management plans. APSAs occurred in areas with significantly lower fishing effort, as indicated by the Mann-Whitney-Wilcoxon test (p < 0.001; Fig. 4a). Over one-third of the APSAs overlapped with existing management areas; however, only 1.6% occurred within areas banning bottom trawling (Fig. 4b). In contrast, 92.6% of APSAs overlapped with ISRAs.

**Fig. 4.**
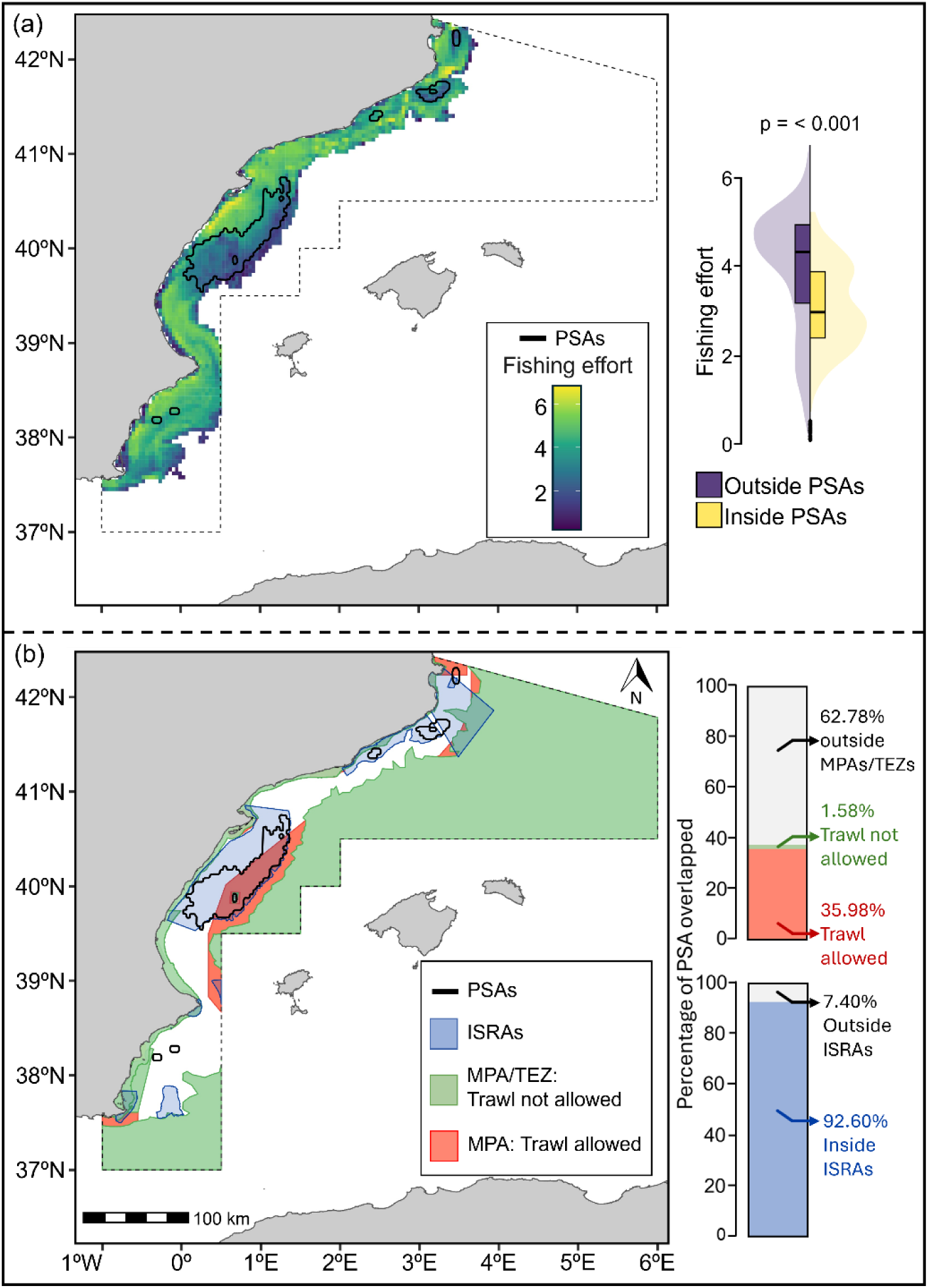
(a) Fishing effort measured as log-transformed (log(x+1)) total trawled hours from 2014 to 2022, with black contour lines delineating Aggregated Potential Spawning Areas (APSAs). The violin plot shows the distribution of fishing effort values inside (yellow) and outside (purple) APSAs; (b) Distribution of Marine Protected Areas (MPAs) and Trawl Exclusion Zones (TEZs) categorised by bottom trawling allowance or ban, along with Important Shark and Ray Areas (ISRAs) as areas of potential conservation priority. Black contour lines delineate APSAs. The upper stacked bar plot shows the percentage of APSAs overlapped by each level of protection against bottom trawling, while the lower one displays the percentage overlapped by ISRAs.

## DISCUSSION

Our findings underscore the urgent need to address bycatch and habitat degradation threats affecting elasmobranch spawning areas. By applying a novel approach to delineate Potential Spawning Areas (PSAs), we identified key environmental and anthropogenic drivers shaping the spatial distribution of egg case bycatch for skates and the smallspotted catshark in the western Mediterranean. Understanding bycatch dynamics for these taxa fills key knowledge gaps regarding fisheries-related impacts on spawning habitats, informing conservation strategies for more vulnerable elasmobranchs. Our findings reveal significant gaps in current spatial management, with a substantial egg case bycatch occurring within PSAs. These results demonstrate the effectiveness of this approach as a tool for identifying critical habitats, providing a robust foundation for prioritizing conservation efforts, including the establishment of targeted Marine Protected Areas (MPAs). Scaling up this approach across other areas and species could further support ecosystem-based management and biodiversity conservation.

### Egg case bycatch

Gaining insight into bycatch rates for elasmobranch egg cases in commercial trawl fisheries is essential for assessing the threats these species face and inform conservation efforts. This study represents the first effort to document such rates, focusing on a trawl fishery operating in the western Mediterranean. Egg case bycatch was recorded in over half of the surveyed tows, with BPUE values approaching 1,000 egg cases per km^2^ for skates and surpassing this threshold for the smallspotted catshark. Approaching this benchmark, often used in fisheries-independent surveys to define high-density areas associated to egg nurseries, underscores an elevated bycatch rate (10, 12, 20).

While egg case abundance for skates of the genera *Raja* and *Bathyraja* typically ranges from 200 to 20,000 per km² in nurseries and surrounding areas (11–13, 15, 16, 31), no bycatch rate estimates have been reported for the smallspotted catshark or other species of the same family. Our observed BPUE values indicate that commercial trawl fisheries intersect high-density areas likely associated to egg nurseries or adjacent habitats (10, 12, 20). This overlap is particularly concerning given the methodological differences between commercial trawl and fisheries-independent surveys, as the latter employs shorter tow distances that optimize the detection of small high-density patches, while the longer tow distances of commercial trawling may dilute evidence of small-sized egg nurseries (12, 16).

#### Static environmental drivers of PSAs

Differences in how skates and the smallspotted catshark respond to the environmental drivers of PSA distribution highlight their species-specific habitat preferences. In the case of the smallspotted catshark, bathymetry was identified as the primary static predictor of bycatch risk, with higher bycatch probabilities in deeper areas of the continental shelf and along the upper slope (100-300 m deep). This range aligns with the species preferred bathymetry and previously documented spawning areas (14, 28, 32). In contrast, skate egg cases were more abundant in the shallower areas (< 100 m), also consistent with known spawning areas (33, 34).

Slope was an important predictor for skate egg case distribution, with higher bycatch probabilities in flat areas (<1°), which is consistent with the preferences of *Bathyraja* skates in the eastern Bering Sea (20). Flat seabeds offer stable environments that minimize disturbances such as sediment resuspension or strong currents - conditions essential for the stability of skate egg cases, which rely solely on adhesion fibres and curved horns for attachment (20, 34). Additionally, seagrass beds or low algal covers commonly associated with flat areas may further stabilize the egg cases and support hatchling development (34). Conversely, smallspotted catshark egg cases exhibited a higher probability of bycatch in gently sloping areas (3°–8°). Catsharks have adapted to these less stable environments by possessing egg cases with tendrils that allow attachment, finding anchorage in habitat-forming invertebrates, such as hydrozoans and soft corals (6, 14). The reliance on biotic structures underscores the ecological significance of these habitat features for egg case stability and survival.

#### Dynamic environmental drivers of PSAs

Sea bottom salinity emerged as a relevant dynamic driver of PSA distribution, with slightly lower salinity relative to the studied range (<38.25 psu) associated with higher probabilities of skate egg case occurrence and increased BPUE for smallspotted catshark egg cases. This finding aligns with prior studies highlighting the role of salinity in defining nursery areas for viviparous sharks, such as school sharks (*Galeorhinus galeus*) favouring high-salinity areas (35) or bull sharks (*Carcharhinus leucas*) preferring low-salinity environments (36), though its role for oviparous elasmobranch species remains unclear.

Sea bottom temperature influenced egg case BPUE for the smallspotted catshark, with higher abundances in cooler areas and seasons, peaking at 13-14°C. This aligns with the species preferred thermal range, associated to the stable conditions of the lower continental shelf and upper slope (37).

Warmer temperatures (15-20°C) have been reported to accelerate embryonic growth and yolk consumption, leading to smaller hatchlings with potentially reduced survival probabilities (38). Such thermal threshold likely explains the seasonal pattern observed, with higher abundances in the cooler seasons (winter and spring), a trend previously reported (39). During cooler periods, PSAs also expand into shallower regions where, despite greater temperature variability, embryo development can still occur, given that it lasts between 4 and 6 months (38). However, rising sea temperatures due to climate change may pose a significant threat to embryo development, particularly in such shallower areas.

#### The feedback loop between fishing pressure and habitat degradation

Trawl fishing effort constituted a key predictor of bycatch risk for both smallspotted catshark and skate egg cases, with lower fishing effort linked to higher BPUE. Only in the case of the catshark, egg case occurrence exhibited greater resilience to moderate fishing pressure but also declined under high pressure. Elevated fishing pressure not only continuously removes specimens and egg cases, but the towed gear also exacerbates habitat degradation, reducing the availability of suitable spawning habitats (5, 6). Moreover, the abrasion further degrades benthic ecosystems by removing biogenic fauna, altering the sediment composition, and reducing ecosystem productivity, availability of organic matter and overall carrying capacity (17, 27). This loss of biogenic structures has been associated to a behavioural shift for smallspotted catsharks, which use plastic debris as oviposition substrates when biological substrates are unavailable, potentially altering the distribution of spawning areas (6). Furthermore, trawling-induced sediment disturbances can lead to the resuspension of fine particles, increasing burial risk (40).

#### Spatial management on APSAs

Estimates of total egg case bycatch indicate that a substantial proportion occurs within PSAs, underscoring the risks posed by trawl fisheries and reinforcing the need to delineate and prioritize these areas for management. By integrating the PSAs predicted for the smallspotted catshark and skates, into multi-species elasmobranch Aggregated Potential Spawning Areas (APSAs), we have identified an area of significant ecological and reproductive importance for oviparous elasmobranchs in the western Mediterranean.

Our approach to defining APSAs is conceptually aligned with criteria used to identify Ecologically and Biologically Significant Marine Areas (EBSAs) or Key Biodiversity Areas (KBAs), emphasizing areas of ecological importance, particularly for the aggregation of critical life-history stages. However, APSAs differ by being rooted in predictive models of bycatch risk, providing a data-driven framework for pinpointing critical habitats. Our multi-species APSAs approach also aligns with ecosystem-based management principles, providing actionable insights to support conservation strategies and inform international policies, such as the EU Maritime Spatial Planning (MSP) Directive, the Marine Strategy Framework Directive (MSFD) or the global “30 by 30” programme under the Kunming-Montreal Global Biodiversity Framework.

Translating these spatial predictions into effective management requires addressing the anthropogenic pressures threatening these habitats, particularly from bottom trawl fisheries. Across the study area, trawl fishing effort is concentrated on the continental shelf and decreases along the slope, leaving limited refugia in highly trawled areas, primarily related to rocky areas, which may also play an important role for skate reproduction (13). A well-designed network of Marine Protected Areas (MPAs) can help mitigate fishing pressure and habitat degradation (41). The highest MPA coverage in the Mediterranean occurs in the western region, with the study area leading in theoretical protection (42), however most MPAs are coastal, leaving significant conservation gaps on the continental shelf and slope (43). Although over a third of the APSAs overlaps with MPAs, only 1.6% lies within areas where bottom trawling is banned, offering limited protection. This underrepresentation of protection beyond coastal areas reflects a general trend across the Mediterranean, as highlighted previously for demersal elasmobranch conservation in the study area (44).

The identified APSAs have the potential to be recognized as Essential Fish Habitats under the GFCM 2030 strategy (23, 24). Furthermore, considering that APSAs are often associated with biogenic habitats formed by hydrozoans and soft corals, recognizing these habitats under the EU Habitats Directive could extend legal protections (6). Notably, almost the entire extent of the APSAs overlaps with ISRAs, established using a biocentric and multi-species framework (28), which further underscores the need for protection. Official endorsement of ISRAs by governmental agencies could play a pivotal role in translating these spatial insights into practical conservation measures promoting the long-term sustainability of these ecologically important areas.

#### Future research

Future studies should focus on identifying specific egg nurseries within the defined APSAs. While our predictions highlight areas suitable for spawning with relatively high BPUE, the definition of egg nurseries requires detecting localised patches of exceptionally high abundances (>1,000 egg cases per km^2^), along with evidence of consistent use over multiple years (9, 10). Incorporating systematic reporting of elasmobranch egg cases into fisheries-independent monitoring surveys in the study area using shorter tow distances could significantly enhance efforts to identify and delineate these critical nursery habitats (9).

Imaging surveys using tools such as towed and drop-down cameras or remotely operated vehicles (ROVs), could also further facilitate the precise delineation of egg nurseries, along with the description of their environmental characteristics, and associated fauna (45, 46). Additionally, unlike trawl surveys, these tools can enable access to hard-substrate areas, which may serve as skate egg nurseries, previously reported in both soft and hard substrate habitats (12, 15, 16, 28). Recent advances in animal tracking technology may also offer promising opportunities to investigate movement ecology in demersal species, providing valuable insights into spawning areas (47).

In addition, further research is needed to assess the population-level impacts of the reported egg case bycatch rates, particularly given the remaining uncertainty surrounding the fate of captured egg cases - whether they can sink, settle, and successfully develop after being discarded. The vulnerability of embryos to bycatch may threat recruitment processes, potentially driving long-term declines in local populations. Evaluating the post-capture viability of the embryos and identifying the environmental and handling factors that influence their survival are critical steps. These findings will be instrumental in improving discarding practices and advancing conservation efforts, as previously suggested for juvenile and adult elasmobranchs (48).

## Conclusions

Elasmobranch conservation is at a pivotal moment, requiring effective measures to protect their critical habitats. We introduce the concept Potential Spawning Areas (PSAs) to define habitats offering optimal conditions for oviposition and identify key reproductive areas for skates and the smallspotted catshark in the western Mediterranean. Our findings reveal a concerning feedback loop between trawl fishing pressure and habitat degradation affecting spawning areas, underscoring the need for targeted conservation measures. Integrating the identified PSAs across species into Aggregated Potential Spawning Areas (APSAs) allows for the recognition of ecologically significant zones and provides a scalable tool for guiding conservation strategies. Moving forward, science-based interventions, such as expanding MPA coverage to include APSAs, enhancing bycatch monitoring, and incorporating predictive models into fisheries management, will be vital for safeguarding these critical habitats. This framework has global applicability, offering a pathway to promote the sustainable management and recovery of elasmobranch populations and their ecosystems.

## MATERIALS AND METHODS

### Study area and data collection

This study was conducted in the Mediterranean waters of the GFCM-designated geographical subarea “GSA06 - Northern Spain”, encompassing bottom trawling grounds at depths from 20 to 700 m. Sampling comprised 194 tows aboard 20 commercial bottom trawlers, conducted as part of the Catalan monitoring survey for commercial fisheries (ICATMAR, available at https://icatmar.cat; 66%) and a demersal elasmobranch monitoring survey (ECEME available at https://eceme.blogs.uv.es; 34%). Seasonal observation campaigns were conducted between December 2020 and June 2022, using a stratified random sampling design proportional to the metiers operating in each port, including: the hake metier on the continental shelf (50-200 m), the Norway lobster metier on the upper slope (200-500 m), and the blue and red shrimp metier on lower slope (500-800 m) (32) (SI Appendix, Fig. S1). The activity was conducted in accordance with the Regulation of the European Parliament and Council on fishing in the GFCM Agreement area, which amends Council Regulation (EC) No. 1967/2006, and did not require ethical review or approval.

Fishing operations were tracked using a GPS (Garmin Ltd., USA), while the SCANMAR system (Scanmar AS, Norway), recorded trawl arrival and retrieval times from the seabed, allowing for the measurement of effective towing track and duration. All operations followed commercial practices and fishing gears consisted of a bottom otter trawl design, with nets averaging a horizontal opening of 24.1 ± 8.5 m across metiers (21.0 ± 7.0 m, 26.9 ± 9.9 m, and 26.3 ± 6.9 m for the hake, Norway lobster and blue and red shrimp metiers, respectively) and a cod-end mesh size of 40 mm, except in Palamós, where 50 mm squared is mandatory only for the blue and red shrimp metier (SI Appendix, Supplementary Fig. S1). On average hauls lasted 3.3 ± 1.5 h at a towing speed of 2.9 ± 0.4 knots, with metier-specific averages of 3.4 ± 1.1 h and 3.2 ± 0.3 knots for hake; 3.3 ± 1.2 h and 2.8 ± 0.3 knots for Norway lobster; and 3.4 ± 2.1 h and 2.6 ± 0.3 knots, for blue and red shrimp.

Elasmobranch egg cases were classified and identified in each tow following (49) and (34). Bycatch per unit effort (BPUE), was calculated as the number of egg cases captured per square kilometre (N/km²). Due to limited species-level identifications of skate egg cases (25.9% of egg cases from 42.0% of tows with occurrences), data were aggregated across *Raja* spp., providing a generalized perspective for modelling bycatch risk.

### Environmental and anthropogenic variables

We evaluated the influence of ten predictor variables, selected based on previous studies, including nine environmental variables (static and dynamic) and one anthropogenic pressure (SI Appendix, Table S6). Variables were sourced from public repositories at the finest spatial resolution available (Table 2; SI Appendix, Fig. S8). All rasters were resampled to a grid resolution of 0.042° using bilinear interpolation. The trawling sampling locations were subsequently aligned both geographically and bathymetrically with the corresponding grid cells to extract data specific to those locations, depths and periods.

### Modelling framework

We utilised Boosted Regression Tree (BRT), a machine-learning approach, to model bycatch risk of egg cases in relation to the predictor variables. BRTs account for non-linear relationships and variable interactions, making them suitable for bycatch risk predictive analyses (50). While collinearity among predictors does not affect BRT predictions, it can influence model interpretation (51). To address this, collinearity was assessed using Spearman correlation (threshold > 0.7; SI Appendix, Fig. S7). Among correlated pairs, slope was retained over roughness, reflecting better the hydrodynamic conditions and habitat stability while being less sensitive to resolution changes. Depth was retained over nutrients (nitrate and phosphate) due to its broader ecological relevance and the limited role of nutrients in the aphotic zone. Fishing effort, which was highly right-skewed, was log-transformed (log(x+1)) prior to analyses.

A delta model framework (hurdle model) was applied to handle zero-inflated data, with separate BRT models for bycatch occurrence (presence/absence, Bernoulli distribution) and abundance (log-transformed BPUE (log(x+1)), Gaussian distribution). These were integrated by multiplying the predicted probability of occurrence by the predicted abundance to produce estimated BPUE values, reflecting both bycatch likelihood and abundance.

Model optimization followed established guidelines (50), testing parameter combinations for boosting iterations, tree complexity, learning rate, and bag fraction (SI Appendix, Tables S2–S5). In BRT, variable selection occurs as the model predominantly disregards non-informative predictors when fitting trees. However, to explicitly exclude unimportant variables, we included an additional variable with random values between 1 and 100 as a benchmark. Predictors with influence exceeding that of the random variable were included in the final models (52). To account for the potential variability associated to each fishing vessel, we used random groups of vessels in a five-fold cross-validation. Selected models in the cross-validation process exceeded 1,000 trees, minimized the deviance while increased the percentage of the deviance explained, and achieved high AUC scores for occurrence predictions. When faced with ties, we prioritized models with higher learning rates, lower tree complexities, and fewer trees to reduce overfitting. The selected models were tested for spatial autocorrelation (SAC) in the residuals based on Moran’s I test (51), with no SAC being detected in any of the models.

To account for model stochasticity and estimate associated uncertainty in predictions, we employed a bootstrap approach, fitting 100 models. For each bootstrap iteration, presence and absence samples were randomly selected with replacement to match the number of original records (53). Uncertainty was quantified using 95% confidence intervals (CI) for variable influence and partial dependence, providing robust estimates of variable importance and response relationships.

#### PSA identification and seasonal persistency assessment

Daily spatial predictions of egg case BPUE for 2021 were generated using the 100 bootstrap models and a 0.042° resolution grid. Medians and 95% CI were calculated from these daily predictions to address model stochasticity and quantify the associated uncertainty. These daily predictions were subsequently used to calculate the seasonal and annual medians, capturing fine-scale temporal variability. Potential Spawning Areas (PSAs) were identified as those grid cells within the 90th percentile of the BPUE predictions, a threshold previously established for defining critical habitats (54). PSAs were calculated for both the seasonal and annual BPUE medians. Seasonal persistence in the spatial distribution of PSAs was assessed as the proportion of seasons in which a grid cell was identified as part of a PSA, providing a measure of their stability. Additionally, seasonal variations in mean BPUE across the study area were assessed using Kruskal-Wallis and Wilcoxon rank-sum tests with Bonferroni corrections.

The overlap between the smallspotted catshark and skates PSAs was quantified using the Intersection over Union (IoU) index. This metric is calculated as the ratio of the intersection area (shared area between the polygons of each taxon) to the union area (total area covered by both sets), expressed as a percentage. To further integrate the spatial extents of both taxa, their PSAs were merged to create multi-species Aggregated Potential Spawning Areas (APSAs), representing regions of shared ecological importance for oviparous elasmobranchs.

#### Anthropogenic pressures and spatial management

An estimation of the total annual bycatch of egg cases was made by integrating our daily predictions of BPUE with daily trawl fishing effort maps obtained from Global Fishing Watch (https://globalfishingwatch.org). Trawl fishing effort, originally measured in hours trawled within a 0.01° grid, was resampled using bilinear interpolation to match the resolution of the BPUE predictions and their associated 95% CI (0.042°). Effort units were then converted from trawling hours to trawled area by multiplying the hours by the product of average trawling speed and net horizontal opening specific to each metier (see “study area and data collection” section), assigned based on depth ranges (32). The total number of bycatch egg cases in each grid cell was calculated as the product of daily BPUE values (N/km²) and the fishing effort (km²). This calculation was repeated for the daily 95% CI values to account for the uncertainty in total bycatch.

To evaluate the intensity of fishing pressure on the delineated areas, trawl fishing effort within and outside APSAs was compared using Mann-Whitney-Wilcoxon tests, following similar approaches in previous studies (54). The overlap of the unified PSA with Marine Protected Areas (MPAs), Trawling Exclusion Zones (TEZs), and Important Shark and Ray Areas (ISRAs) was analysed using the IoU index to assess the level of protection offered by existing and proposed protected areas. MPA boundaries were sourced from ProtectedSeas Navigator (https://navigatormap.org) while regulatory measures on trawling fisheries were derived from the management plans of each MPA. TEZs, defined by the national fisheries regulatory plan ([Orden APA/1074/2007](https://www.boe.es/buscar/doc.php?id=BOE-A-2007-8559)), restrict bottom trawling to depths exceeding 50 meters, with an exception applying to the continental shelf extension generated by the Ebro Delta (40° 43.2’N to 39° 44.4’N) where trawling is prohibited within 3 nautical miles of the coast. ISRAs, designated by (55), identify important areas for elasmobranchs to inform decision-making but currently lack management plans.

## Data and code availability

All analyses were performed with R 4.4.1 (56). The R code used on these analyses, along with the bycatch data along and the resulting APSA polygons, are available in Zenodo public repositories: https://doi.org/10.5281/zenodo.14847186, and https://doi.org/10.5281/zenodo.14847319, respectively.

## Acknowledgments

We thank the fisheries associations from Catalonia, El Grao de Castellón, Cullera, Jávea, Calpe and Cartagena and particularly to the skippers and the crew from the F/Vs participating in the data collection. We thank the technicians of the Institut Català de Recerca per a la Governança del Mar (ICATMAR), who helped with the data collection. We also thank the Conselleria de Agricultura, Desarrollo Rural, Emergencia Climática y Transición Ecológica of the Generalitat Valenciana, the Consejería de Agua, Agricultura, Ganadería, Pesca y Medio Ambiente de la Región de Murcia and the Direcció General de Política Marítima i Pesca Sostenible de la Generalitat de Catalunya (Spain). This research was supported by the Biodiversity Foundation of the Ministry for the Ecological Transition and Demographic Challenge under the project ECEME (CA_BM_2019). DR-G was supported by a FPU grant of the Spanish Ministry of Universities (MIU). DM acknowledges support from the CIDEGENT program of the Generalitat Valenciana (CIDEGENT/2021/058).

## Author Contributions

D.R-G, C.B. and D.M. designed research; D.R-G, C.B., A.I.C., J.A.R., and D.M. performed research; D.R-G, and D.M. analysed data; and D.R-G, C.B., A.I.C., J.A.R., and D.M. wrote the paper.

## Notes

https://doi.org/10.5281/zenodo.14847186

https://doi.org/10.5281/zenodo.14847319

### Competing Interest Statement

The authors have declared no competing interest.

