## Supplementary material for "Predicting Potential Spawning Areas: a novel framework for elasmobranch conservation and spatial management": SI Appendix

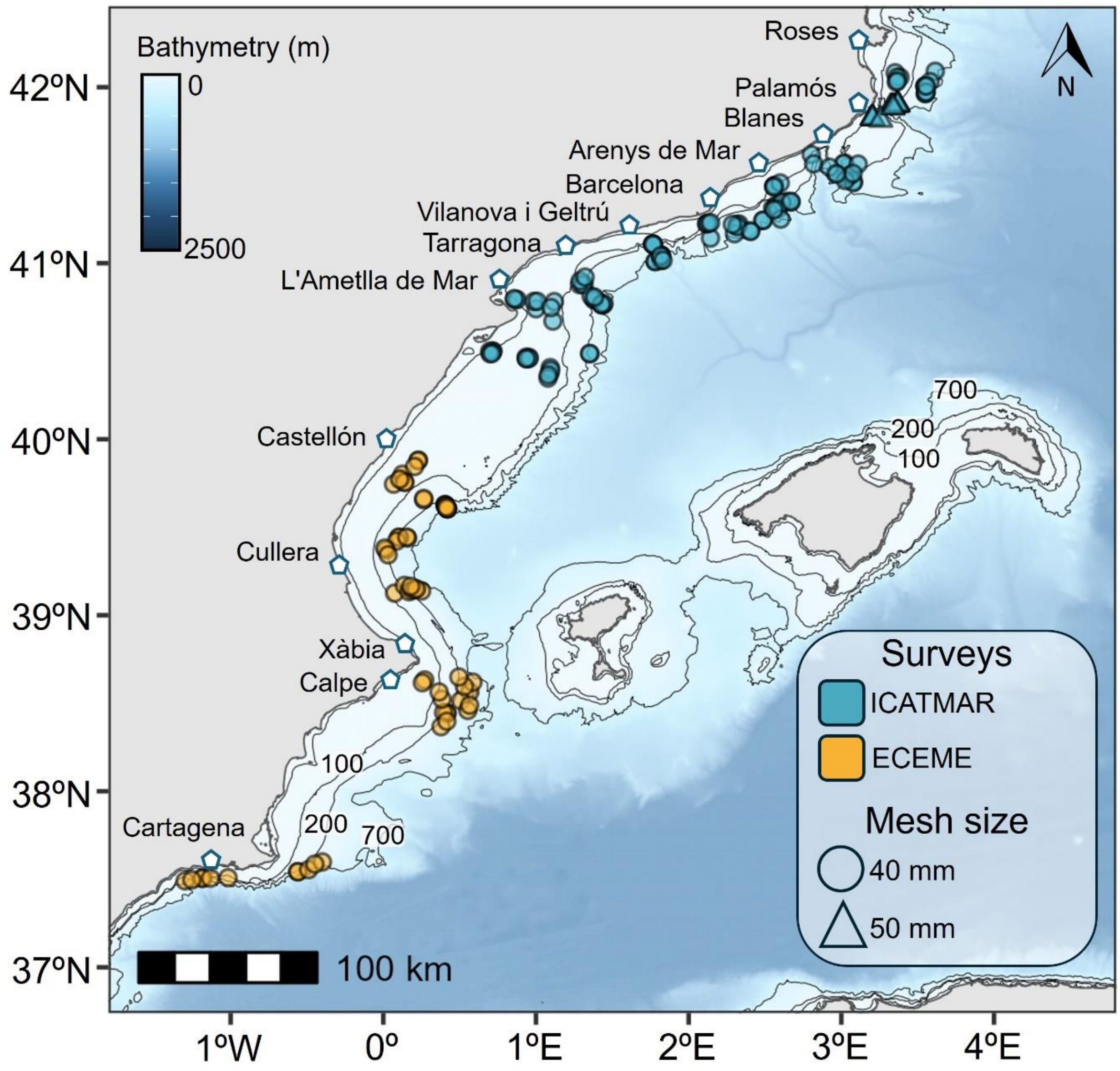


**Fig. S1.** Map of the study area illustrating the locations of the 194 tows analysed. The colour coding indicates the survey from which the data was obtained, while different shapes denote the used cod-end mesh size. Studied ports are highlighted on the map using pentagon symbols.

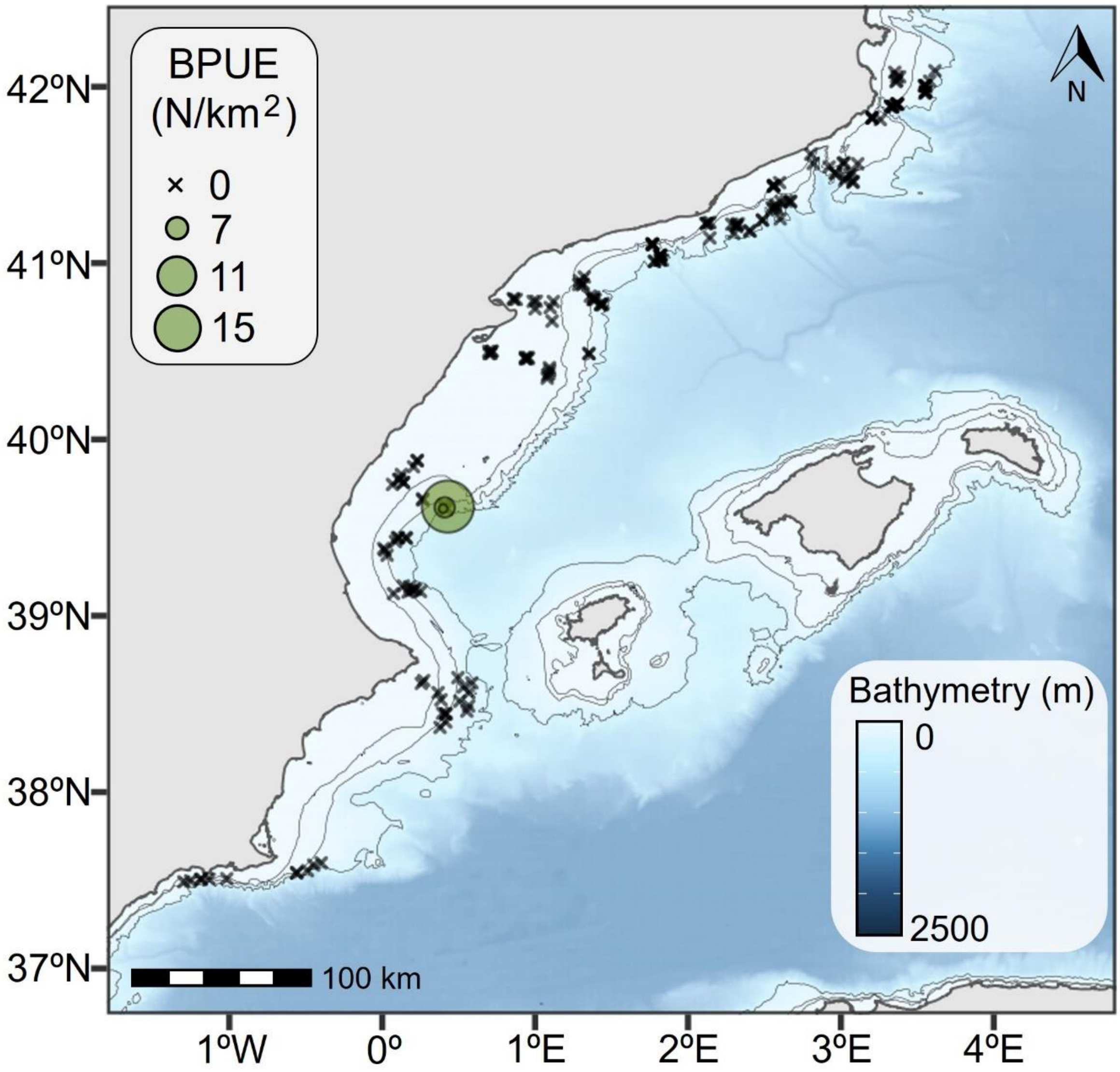


**Fig. S2.** Distribution of longnosed skate (*Dipturus oxyrinchus*) egg case BPUE. Circles represent egg case presence, with size proportional to BPUE, while crosses denote their absence at sampling location.

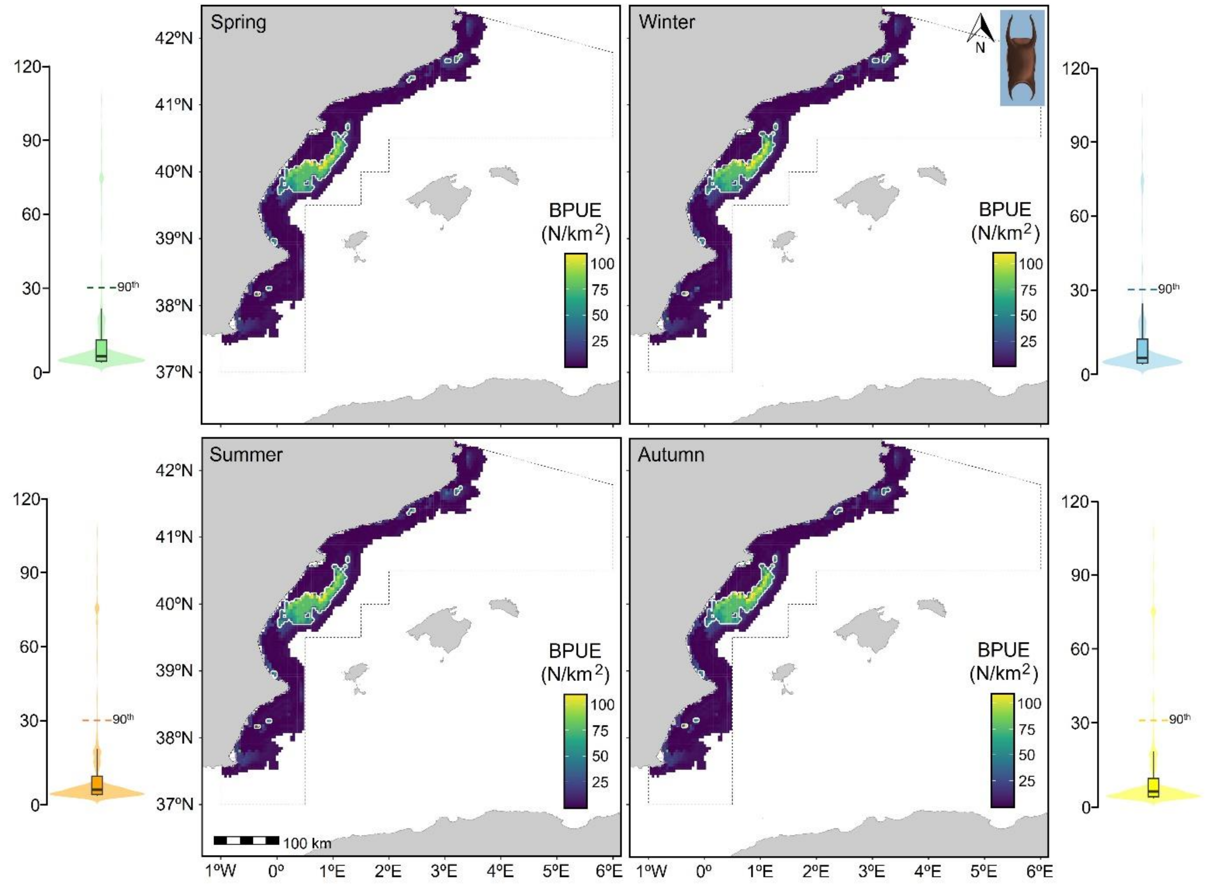


**Fig. S3**. Seasonal median BPUE predictions, based on the daily median of bootstrap models (n = 100) for skates (*Raja* spp.). Violin plot shows the distribution of BPUE values for each season and the threshold of the 90th percentile.

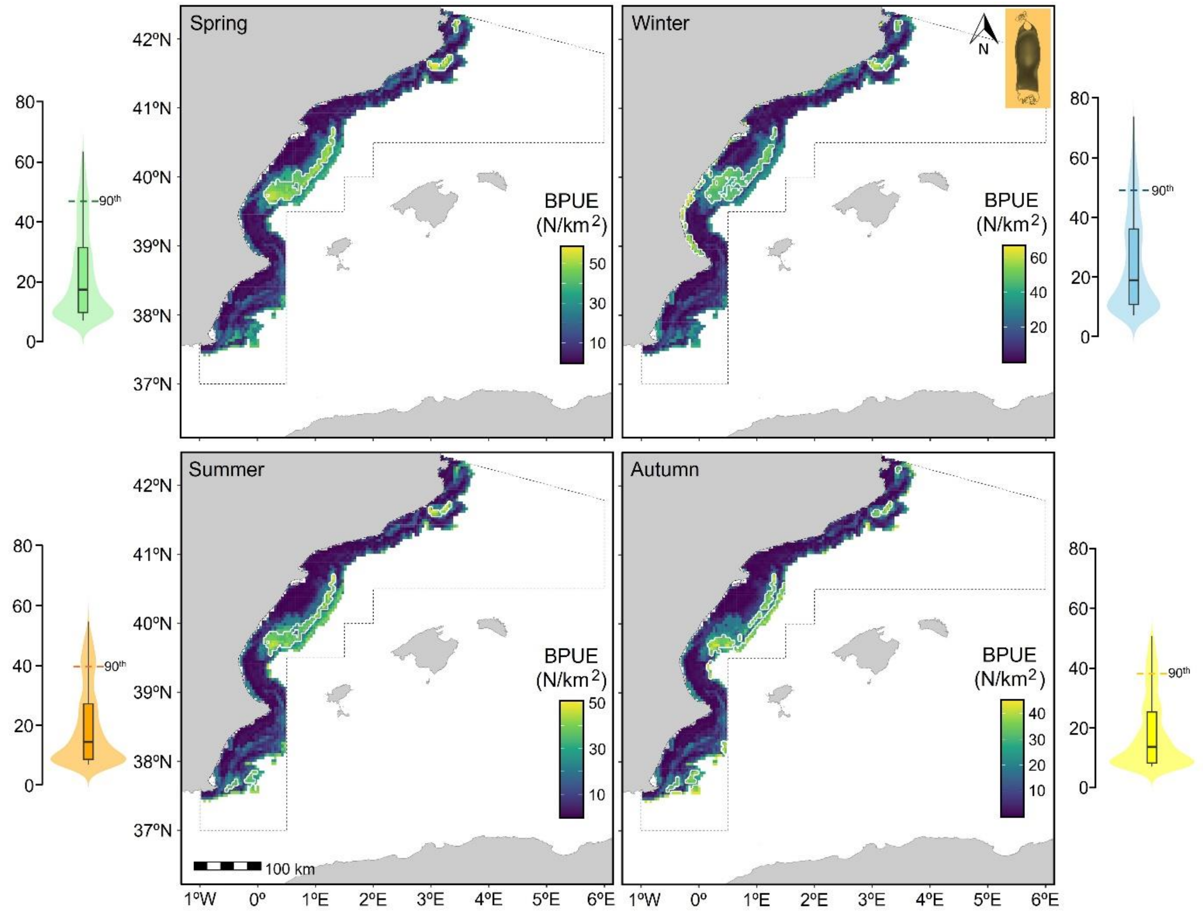


**Fig. S4**. Seasonal median BPUE predictions, based on the daily median of bootstrap models (n = 100) for the small-spotted catshark (*Scyliorhinus canicula*). Violin plot shows the distribution of BPUE values for each season and the threshold of the 90th percentile.

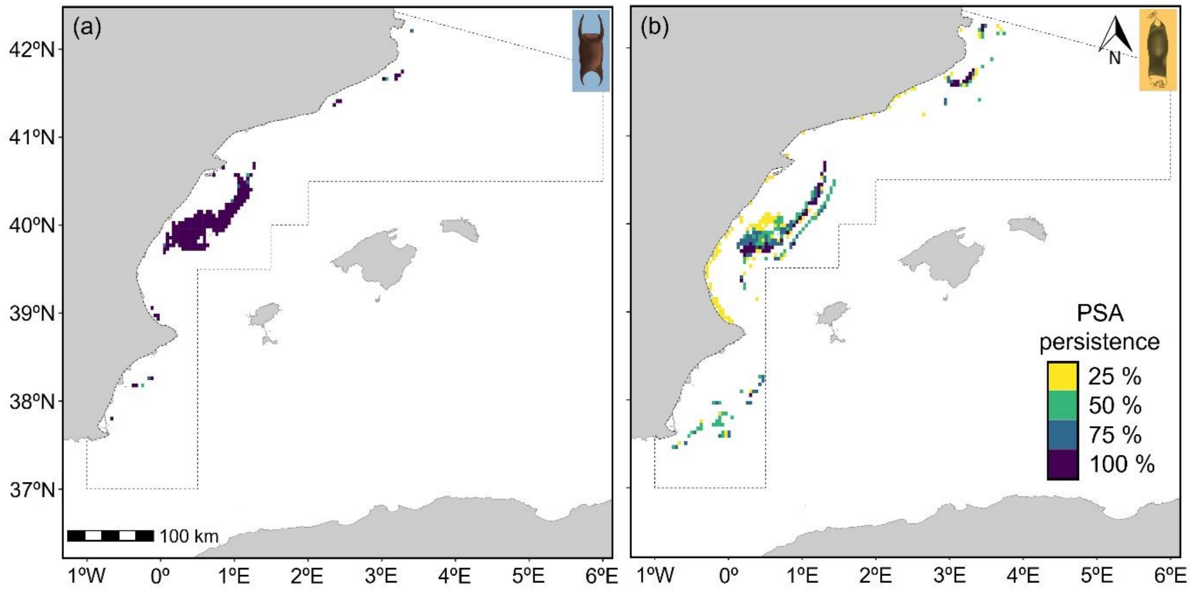


**Fig. S5**. Persistence of PSAs across seasons for (a) skates, and (b) smallspotted catshark egg cases.

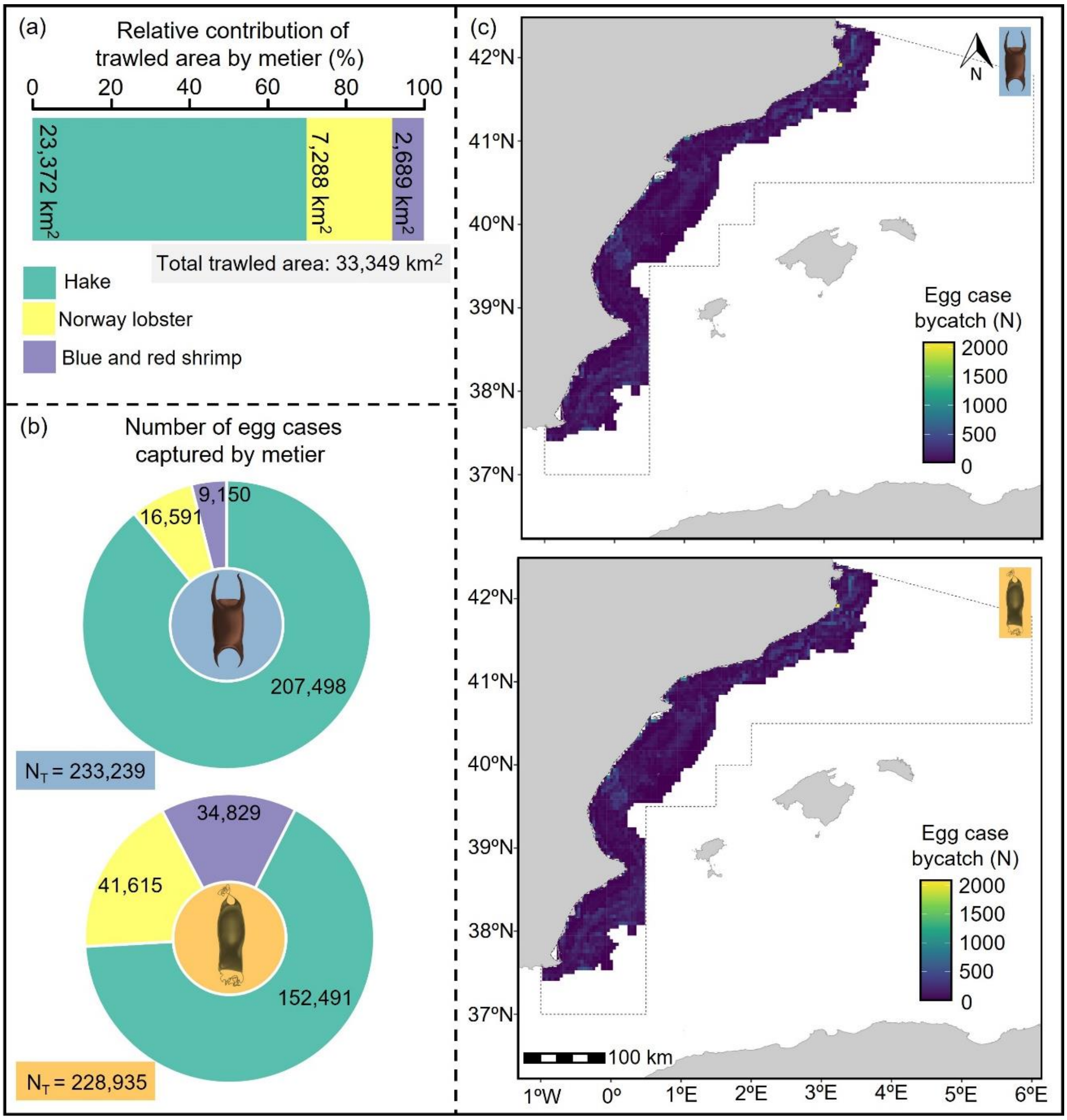


**Fig. S6**. Estimated annual total bycatch figures for 2021. (a) Total area trawled by each metier within the study area; (b) total number of egg cases for skates and the small-spotted catshark by each metier; and (c) spatial distribution of estimated annual total egg case bycatch.

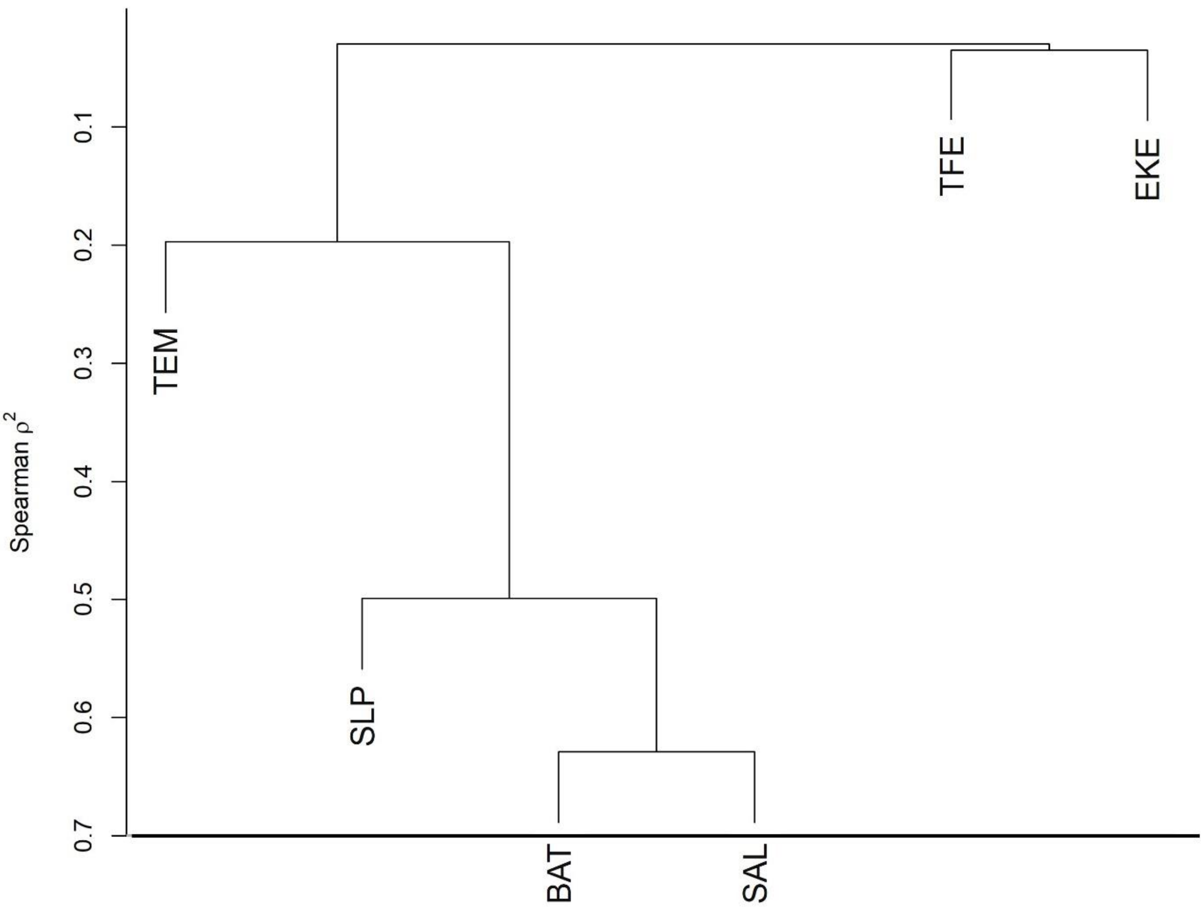


**Fig. S7**. Hierarchical cluster on exploratory variables based on squared Spearman correlation. The grey line represents the 0.7 threshold used for assessing collinearity. Predictor acronyms, definitions and units are described in Table 2.

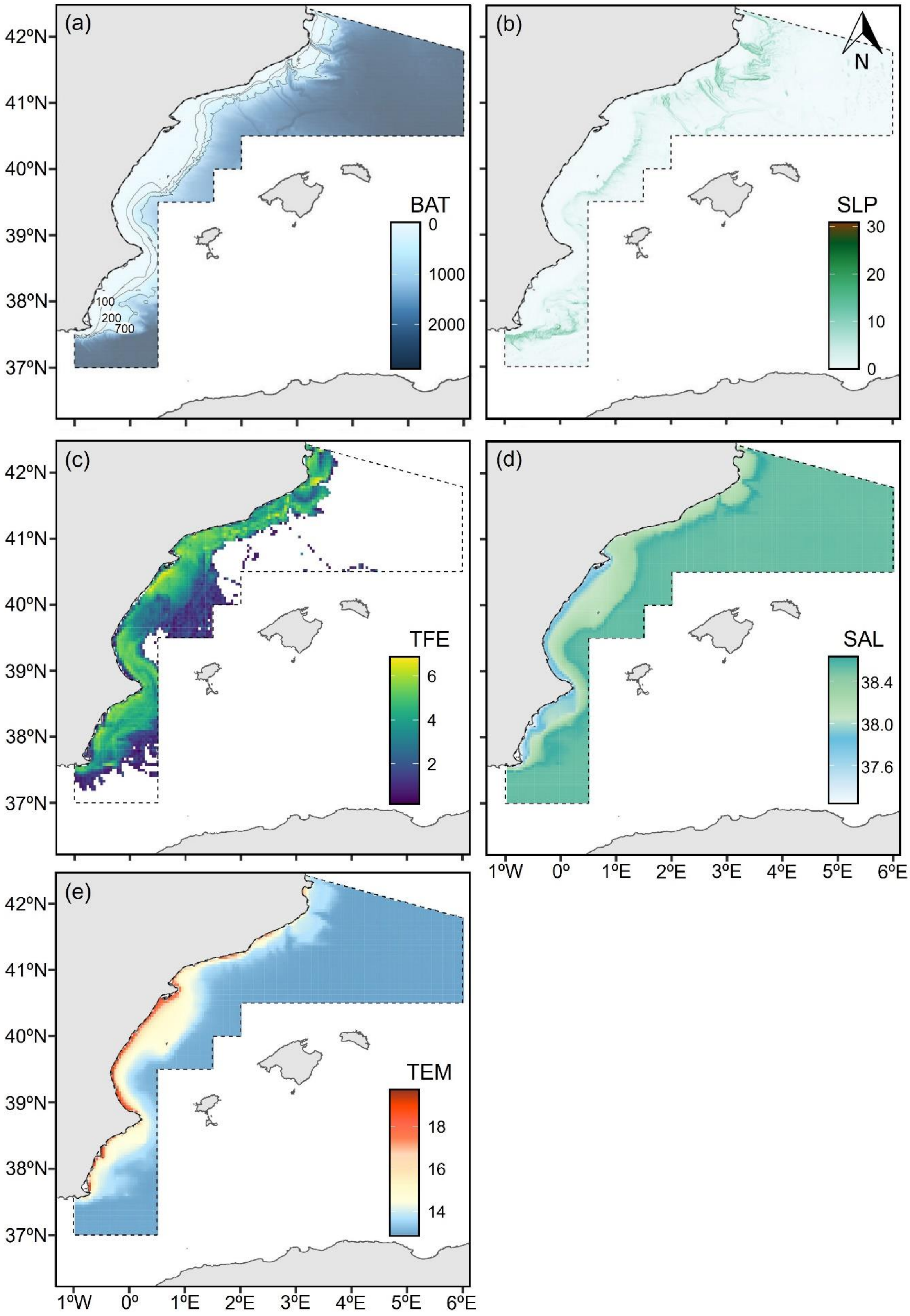


**Fig. S8.** Environmental and human pressure drivers kept in the models to predict the distribution of egg cases, including: (a) bathymetry (BAT); (b) slope (SLP); (c) log-transformed trawl fishing effort (log(x+1)) (TFE); (d) sea bottom salinity (SAL); and (e) sea bottom temperature (TEM). Predictor descriptions and units are described in Table 2.

**Table S1**. List of skate species of genus *Raja* for which egg cases were captured, with their conservation status based on the regional assessment of the Mediterranean IUCN Red List. Reported values include the total number of egg cases captured ($N_T$), the maximum BPUE (N/km$^2$) with the corresponding number of egg cases (n) in a single tow, the mean BPUE (N/km$^2$), the frequency of occurrence (Freq.) across all surveyed tows, and the depth range in which they occurred.

| **Species** | **IUCN status** | **$N_T$** | **maximum** | | **Mean BPUE** | **Frequency (%)** | **Depth range (m)** |
|---|---|---|---|---|---|---|---|
| | | | **BPUE** | **n** | | | |
| *Raja asterias* | NT | 185 | 331.4 | 95 | 10.2 ± 44.4 | 87.3 | 55-595 |
| *Raja polystigma* | LC | 13 | 16.2 | 4 | 0.2 ± 2.0 | 6.1 | 55-176 |
| *Raja clavata* | NT | 10 | 7.8 | 3 | 0.4 ± 1.5 | 4.7 | 65-320 |
| *Raja miraletus* | LC | 3 | 18.4 | 1 | 0.7 ± 2.8 | 1.4 | 65-69 |
| *Raja radula* | EN | 1 | 5.2 | 1 | 0.1 ± 0.8 | 0.5 | 73 |

**Table S2.** Fitted combinations of boosted regression tree parameters used for optimizing the model to predict bycatch occurrence (presence/absence) of skates (*Raja* spp.) using a bernoulli distribution. The results are ordered by cross-validated deviance, from smallest to largest. Parameters selected for analysis are highlighted with grey shading.

| Learning rate | Tree complexity | Bag fraction | Number of trees | Cross-validation AUC | Cross-validation deviance | Cross-validation deviance explained (%) |
|---|---|---|---|---|---|---|
| 0.0010 | 4 | 0.8 | 2150 | 0.797 | 0.872 | 29.502 |
| 0.0005 | 4 | 0.8 | 4450 | 0.799 | 0.873 | 29.448 |
| 0.0050 | 4 | 0.8 | 450 | 0.797 | 0.873 | 29.409 |
| 0.0005 | 5 | 0.8 | 4400 | 0.795 | 0.875 | 29.289 |
| 0.0050 | 5 | 0.8 | 450 | 0.792 | 0.876 | 29.223 |
| 0.0050 | 3 | 0.8 | 500 | 0.796 | 0.876 | 29.206 |
| 0.0010 | 5 | 0.8 | 2200 | 0.793 | 0.876 | 29.148 |
| 0.0005 | 3 | 0.8 | 4950 | 0.795 | 0.877 | 29.127 |
| 0.0010 | 3 | 0.8 | 2650 | 0.795 | 0.879 | 28.927 |
| 0.0010 | 4 | 0.7 | 2450 | 0.794 | 0.880 | 28.899 |
| 0.0005 | 4 | 0.7 | 4850 | 0.792 | 0.881 | 28.793 |
| 0.0050 | 4 | 0.7 | 450 | 0.792 | 0.882 | 28.720 |
| 0.0050 | 5 | 0.7 | 450 | 0.791 | 0.882 | 28.689 |
| 0.0010 | 4 | 0.6 | 2550 | 0.787 | 0.883 | 28.633 |
| 0.0005 | 3 | 0.7 | 4700 | 0.794 | 0.883 | 28.590 |
| 0.0010 | 5 | 0.7 | 2450 | 0.789 | 0.884 | 28.563 |
| 0.0005 | 4 | 0.6 | 4950 | 0.788 | 0.885 | 28.449 |
| 0.0005 | 5 | 0.7 | 4650 | 0.786 | 0.885 | 28.449 |
| 0.0050 | 3 | 0.7 | 500 | 0.791 | 0.885 | 28.433 |
| 0.0005 | 5 | 0.6 | 4700 | 0.785 | 0.886 | 28.398 |
| 0.0005 | 3 | 0.6 | 5500 | 0.792 | 0.886 | 28.352 |
| 0.0050 | 4 | 0.6 | 600 | 0.792 | 0.887 | 28.279 |
| 0.0010 | 3 | 0.7 | 2800 | 0.790 | 0.887 | 28.279 |
| 0.0010 | 5 | 0.6 | 2450 | 0.789 | 0.887 | 28.266 |
| 0.0010 | 3 | 0.6 | 2900 | 0.789 | 0.888 | 28.217 |
| 0.0050 | 3 | 0.6 | 550 | 0.787 | 0.889 | 28.163 |
| 0.0050 | 5 | 0.6 | 400 | 0.788 | 0.892 | 27.874 |
| 0.0001 | 4 | 0.8 | 10000 | 0.797 | 0.925 | 25.258 |
| 0.0001 | 5 | 0.8 | 10000 | 0.789 | 0.927 | 25.095 |
| 0.0001 | 3 | 0.8 | 10000 | 0.803 | 0.927 | 25.024 |
| 0.0001 | 4 | 0.7 | 10000 | 0.790 | 0.934 | 24.472 |
| 0.0001 | 5 | 0.7 | 10000 | 0.788 | 0.936 | 24.375 |
| 0.0001 | 3 | 0.7 | 10000 | 0.796 | 0.937 | 24.236 |
| 0.0001 | 4 | 0.6 | 10000 | 0.789 | 0.942 | 23.865 |
| 0.0001 | 5 | 0.6 | 10000 | 0.785 | 0.943 | 23.807 |
| 0.0001 | 3 | 0.6 | 10000 | 0.788 | 0.946 | 23.517 |

**Table S3.** Fitted combinations of boosted regression tree parameters used for optimizing the model to predict BPUE of skates (*Raja* spp.) using a gaussian distribution. Results are ordered from smallest to largest cross-validated deviance. Selected parameters for analysis are marked with a grey shading.

| Learning rate | Tree complexity | Bag fraction | Number of trees | Cross-validation deviance | Cross-validation deviance explained (%) |
|---|---|---|---|---|---|
| 0.0010 | 4 | 0.8 | 2850 | 1.395 | 46.178 |
| 0.0010 | 3 | 0.8 | 3000 | 1.397 | 46.086 |
| 0.0005 | 3 | 0.8 | 5700 | 1.397 | 46.079 |
| 0.0050 | 5 | 0.8 | 550 | 1.398 | 46.050 |
| 0.0005 | 4 | 0.8 | 5650 | 1.399 | 46.031 |
| 0.0005 | 5 | 0.8 | 5850 | 1.399 | 46.014 |
| 0.0050 | 4 | 0.8 | 550 | 1.399 | 45.998 |
| 0.0050 | 3 | 0.8 | 550 | 1.400 | 45.964 |
| 0.0050 | 3 | 0.7 | 600 | 1.401 | 45.931 |
| 0.0005 | 3 | 0.7 | 6250 | 1.401 | 45.917 |
| 0.0050 | 5 | 0.7 | 700 | 1.402 | 45.897 |
| 0.0050 | 4 | 0.7 | 750 | 1.403 | 45.867 |
| 0.0050 | 3 | 0.6 | 500 | 1.403 | 45.853 |
| 0.0010 | 5 | 0.8 | 2600 | 1.404 | 45.836 |
| 0.0010 | 5 | 0.7 | 2800 | 1.404 | 45.832 |
| 0.0005 | 5 | 0.7 | 6100 | 1.404 | 45.825 |
| 0.0010 | 4 | 0.7 | 3050 | 1.404 | 45.803 |
| 0.0050 | 4 | 0.6 | 500 | 1.405 | 45.773 |
| 0.0010 | 3 | 0.7 | 3050 | 1.406 | 45.730 |
| 0.0010 | 4 | 0.6 | 2550 | 1.408 | 45.667 |
| 0.0010 | 3 | 0.6 | 2700 | 1.409 | 45.637 |
| 0.0005 | 5 | 0.6 | 5200 | 1.410 | 45.582 |
| 0.0050 | 5 | 0.6 | 500 | 1.412 | 45.517 |
| 0.0005 | 4 | 0.6 | 4900 | 1.414 | 45.442 |
| 0.0005 | 3 | 0.6 | 5100 | 1.414 | 45.436 |
| 0.0005 | 4 | 0.7 | 5700 | 1.415 | 45.414 |
| 0.0010 | 5 | 0.6 | 2700 | 1.417 | 45.313 |
| 0.0001 | 5 | 0.8 | 10000 | 1.577 | 39.158 |
| 0.0001 | 4 | 0.8 | 10000 | 1.577 | 39.132 |
| 0.0001 | 3 | 0.8 | 10000 | 1.579 | 39.067 |
| 0.0001 | 3 | 0.6 | 10000 | 1.592 | 38.577 |
| 0.0001 | 4 | 0.6 | 10000 | 1.592 | 38.554 |
| 0.0001 | 5 | 0.6 | 10000 | 1.594 | 38.494 |
| 0.0001 | 4 | 0.7 | 10000 | 1.595 | 38.450 |
| 0.0001 | 5 | 0.7 | 10000 | 1.596 | 38.415 |
| 0.0001 | 3 | 0.7 | 10000 | 1.597 | 38.382 |

**Table S4.** Fitted combinations of boosted regression tree parameters used for optimizing the model to predict bycatch occurrence (presence/absence) of the small-spotted catshark (*Scyliorhinus canicula*) using a bernoulli distribution. The results are ordered by cross-validated deviance, from smallest to largest. Parameters selected for analysis are highlighted with grey shading.

| Learning rate | Tree complexity | Bag fraction | Number of trees | Cross-validation AUC | Cross-validation deviance | Cross-validation deviance explained (%) |
|---|---|---|---|---|---|---|
| 0.001 | 5 | 0.8 | 4250 | 0.795 | 1.110 | 19.782 |
| 0.001 | 4 | 0.8 | 5050 | 0.790 | 1.110 | 19.779 |
| 0.0005 | 4 | 0.8 | 9650 | 0.792 | 1.110 | 19.765 |
| 0.005 | 4 | 0.8 | 1000 | 0.789 | 1.112 | 19.666 |
| 0.0005 | 5 | 0.8 | 8350 | 0.792 | 1.112 | 19.638 |
| 0.001 | 3 | 0.8 | 7000 | 0.792 | 1.113 | 19.540 |
| 0.005 | 3 | 0.8 | 1400 | 0.793 | 1.115 | 19.420 |
| 0.005 | 5 | 0.8 | 850 | 0.788 | 1.118 | 19.193 |
| 0.005 | 4 | 0.7 | 850 | 0.787 | 1.119 | 19.109 |
| 0.0005 | 5 | 0.7 | 7400 | 0.788 | 1.120 | 19.064 |
| 0.005 | 5 | 0.7 | 800 | 0.785 | 1.121 | 19.007 |
| 0.001 | 4 | 0.7 | 4250 | 0.791 | 1.122 | 18.937 |
| 0.001 | 5 | 0.7 | 3900 | 0.788 | 1.122 | 18.929 |
| 0.0005 | 3 | 0.8 | 10000 | 0.787 | 1.123 | 18.848 |
| 0.0005 | 4 | 0.7 | 7950 | 0.785 | 1.126 | 18.640 |
| 0.005 | 5 | 0.6 | 750 | 0.786 | 1.127 | 18.565 |
| 0.005 | 3 | 0.7 | 1250 | 0.787 | 1.129 | 18.420 |
| 0.0005 | 3 | 0.7 | 10000 | 0.784 | 1.131 | 18.265 |
| 0.001 | 5 | 0.6 | 3700 | 0.790 | 1.131 | 18.245 |
| 0.001 | 3 | 0.7 | 5500 | 0.787 | 1.133 | 18.131 |
| 0.005 | 4 | 0.6 | 800 | 0.783 | 1.133 | 18.113 |
| 0.0005 | 5 | 0.6 | 6500 | 0.780 | 1.134 | 18.010 |
| 0.0005 | 4 | 0.6 | 7550 | 0.779 | 1.135 | 17.950 |
| 0.001 | 4 | 0.6 | 4050 | 0.783 | 1.138 | 17.763 |
| 0.001 | 3 | 0.6 | 5200 | 0.783 | 1.144 | 17.313 |
| 0.0005 | 3 | 0.6 | 8150 | 0.776 | 1.150 | 16.865 |
| 0.005 | 3 | 0.6 | 1100 | 0.777 | 1.151 | 16.782 |
| 0.0001 | 5 | 0.7 | 10000 | 0.753 | 1.208 | 12.663 |
| 0.0001 | 5 | 0.8 | 10000 | 0.746 | 1.210 | 12.575 |
| 0.0001 | 5 | 0.6 | 10000 | 0.759 | 1.213 | 12.306 |
| 0.0001 | 4 | 0.7 | 10000 | 0.755 | 1.220 | 11.824 |
| 0.0001 | 4 | 0.8 | 10000 | 0.747 | 1.222 | 11.679 |
| 0.0001 | 4 | 0.6 | 10000 | 0.753 | 1.223 | 11.629 |
| 0.0001 | 3 | 0.8 | 10000 | 0.750 | 1.237 | 10.569 |
| 0.0001 | 3 | 0.7 | 10000 | 0.748 | 1.238 | 10.560 |
| 0.0001 | 3 | 0.6 | 10000 | 0.748 | 1.239 | 10.451 |

**Table S5.** Fitted combinations of boosted regression tree (BRT) parameters used for optimizing the model to predict bycatch predict BPUE of the small-spotted catshark (*Scyliorhinus canicula*) using a gaussian distribution. The results are ordered by cross-validated deviance, from smallest to largest. Parameters selected for analysis are highlighted with grey shading.

| Learning rate | Tree complexity | Bag fraction | Number of trees | Cross-validation deviance | Cross-validation deviance explained (%) |
|---|---|---|---|---|---|
| 0.005 | 3 | 0.6 | 250 | 1.394 | 17.274 |
| 0.005 | 4 | 0.6 | 250 | 1.402 | 16.774 |
| 0.001 | 5 | 0.6 | 1350 | 1.402 | 16.766 |
| 0.001 | 3 | 0.6 | 1400 | 1.404 | 16.678 |
| 0.0005 | 3 | 0.6 | 2800 | 1.407 | 16.491 |
| 0.0005 | 5 | 0.7 | 2550 | 1.407 | 16.457 |
| 0.001 | 4 | 0.7 | 1300 | 1.408 | 16.436 |
| 0.005 | 4 | 0.8 | 250 | 1.409 | 16.366 |
| 0.0005 | 4 | 0.7 | 2750 | 1.409 | 16.364 |
| 0.0005 | 4 | 0.6 | 2950 | 1.409 | 16.355 |
| 0.0005 | 5 | 0.6 | 3000 | 1.409 | 16.340 |
| 0.001 | 5 | 0.7 | 1450 | 1.411 | 16.218 |
| 0.001 | 4 | 0.6 | 1300 | 1.412 | 16.184 |
| 0.005 | 4 | 0.7 | 250 | 1.413 | 16.145 |
| 0.005 | 3 | 0.7 | 250 | 1.413 | 16.124 |
| 0.0005 | 4 | 0.8 | 2600 | 1.414 | 16.083 |
| 0.005 | 5 | 0.6 | 250 | 1.415 | 16.032 |
| 0.001 | 4 | 0.8 | 1200 | 1.415 | 15.993 |
| 0.0005 | 3 | 0.7 | 2750 | 1.415 | 15.984 |
| 0.001 | 5 | 0.8 | 1250 | 1.416 | 15.957 |
| 0.001 | 3 | 0.7 | 1500 | 1.416 | 15.924 |
| 0.001 | 3 | 0.8 | 1300 | 1.417 | 15.904 |
| 0.0001 | 5 | 0.7 | 10000 | 1.417 | 15.879 |
| 0.005 | 5 | 0.8 | 300 | 1.418 | 15.854 |
| 0.005 | 3 | 0.8 | 250 | 1.418 | 15.852 |
| 0.0001 | 4 | 0.7 | 10000 | 1.418 | 15.852 |
| 0.0001 | 5 | 0.6 | 10000 | 1.418 | 15.822 |
| 0.0005 | 3 | 0.8 | 2700 | 1.418 | 15.822 |
| 0.0001 | 4 | 0.6 | 10000 | 1.419 | 15.797 |
| 0.0001 | 3 | 0.6 | 10000 | 1.419 | 15.767 |
| 0.0005 | 5 | 0.8 | 2750 | 1.419 | 15.753 |
| 0.0001 | 4 | 0.8 | 10000 | 1.420 | 15.725 |
| 0.0001 | 5 | 0.8 | 10000 | 1.421 | 15.667 |
| 0.005 | 5 | 0.7 | 250 | 1.422 | 15.617 |
| 0.0001 | 3 | 0.7 | 10000 | 1.424 | 15.485 |
| 0.0001 | 3 | 0.8 | 10000 | 1.426 | 15.349 |

**Table S6.** Relevance of predictors considered to assess egg case bycatch distribution.

| Predictor | Relation to response variable |
|---|---|
| Depth | Shapes species niches through adaptations to varying environmental and biological factors, such as light, pressure, and nutrients, which influence community composition along bathymetric gradients (1, 2). |
| Slope | Reflects the steepness and directional elevation changes of the seabed, which influence current velocity and direction. These factors are linked to habitat stability and may importantly affect egg case attachment and embryo development (3). |
| Roughness | Measures terrain variability by quantifying how uneven or rugged the surface is within a given area, regardless of direction. It underscores habitat complexity, which can influence habitat preferences (4). |
| Substrate | Elasmobranch spawning areas have been observed to occur preferably over different seabed substrates from sandy to rocky areas depending on the species (5–7). The substrate physical composition may therefore influence the distribution of ovipositing areas. However, it must be considered that trawl fisheries only operate over soft bottoms, thus, our dataset can only test the influence of the different soft substrates according to folk categorization (sandy, muddy and mixed sediment bottoms) (8). |
| Trawling fishing effort | Chronically trawled areas experience significant decreases in organic matter, entailing important habitat degradation and loss (9), impacting elasmobranchs that depend on these environments for food, shelter or reproduction, including spawning areas (10, 11). |
| Sea bottom temperature | Most species are narrowly thermally adapted and bottom temperature can influence importantly the development of the embryos (12). |
| Sea bottom salinity | Previous studies suggest that slight changes can play a crucial role in determining nursery habitats for elasmobranch species (13). |
| Nitrate concentration in the sea bottom | Indicator of primary production in shallow bottoms, where they might influence elasmobranch species distribution (14, 15); however, its role in deepwater environments might be limited by light levels impeding their usage in primary production (16). |
| Phosphate concentration in the sea bottom | Indicator of primary production in shallow bottoms, where they might influence elasmobranch species distribution (14, 15); however, its role in deepwater environments might be limited by light levels impeding their usage in primary production (16). |
| Eddy kinetic energy in the sea bottom | A measure of turbulent mixing and variability in current speed, can also influence nutrient distribution and habitat conditions, potentially affecting the distribution of marine species (17). |
